# An Inherent Structural Difference Between Serine and Threonine Phosphorylation: Phosphothreonine Prefers an Ordered, Compact, Cyclic Conformation

**DOI:** 10.1101/2020.02.29.971382

**Authors:** Anil K. Pandey, Himal K. Ganguly, Sudipta Kumar Sinha, Kelly E. Daniels, Glenn P. A. Yap, Sandeep Patel, Neal J. Zondlo

**Author notes:** These authors contributed equally (Anil K. Pandey, Himal K. Ganguly). Department of Chemistry, Indian Institute of Technology Ropar, Nangal Road 140001 India (Sudipta Kumar Sinha).

## Abstract

Phosphorylation and dephosphorylation of proteins by kinases and phosphatases are central to cellular responses and function. The structural effects of serine and threonine phosphorylation were examined in peptides and in proteins, by circular dichroism, NMR spectroscopy, bioinformatics analysis of the PDB, small-molecule X-ray crystallography, and computational investigations. Phosphorylation of both serine and threonine residues induces substantial conformational restriction in their physiologically more important dianionic forms. Threonine exhibits a particularly strong disorder-to-order transition upon phosphorylation, with dianionic phosphothreonine preferentially adopting a cyclic conformation with restricted *ϕ* (*ϕ* ∼ –60°) stabilized by three noncovalent interactions: a strong intraresidue phosphate-amide hydrogen bond, an n→π* interaction between consecutive carbonyls, and an n→σ* interaction between the phosphate Oγ lone pair and the antibonding orbital of C–Hβ that restricts the *χ*_2_ side chain conformation. Proline is unique among the canonical amino acids for its covalent cyclization on the backbone. Phosphothreonine can mimic proline’s backbone cyclization via noncovalent interactions. The preferred torsions of dianionic phosphothreonine are *ϕ,ψ* = polyproline helix or α-helix (*ϕ* ∼ –60°); *χ*_1_ = *g*^−^; *χ*_2_ = eclipsed C–H/O–P bonds. This structural signature is observed in diverse proteins, including the activation loops of protein kinases and protein-protein interactions. In total, these results suggest a structural basis for the differential use and evolution of threonine versus serine phosphorylation sites in proteins, with serine phosphorylation typically inducing smaller, rheostat-like changes, versus threonine phosphorylation promoting larger, step function-like switches, in proteins.

## Introduction

Phosphorylation of serine (Ser), threonine (Thr), and tyrosine (Tyr) hydroxyl groups in proteins is a predominant mechanism of intracellular signal transduction and of achieving intracellular responses to extracellular signals.^1^ Over 90% of phosphorylated residues identified via phosphoproteomics are phosphoserine (pSer) or phosphothreonine (pThr) residues, with the significant majority of these phosphorylation sites being pSer residues.^2^ The preference of phosphorylation on Ser over Thr is observed in most proteins, via biases at the substrate level both of kinases (faster phosphorylation of Ser than Thr) and phosphatases (faster dephosphorylation at Thr than Ser). These general substrate preferences combine to result in a significantly higher fraction of pSer than pThr residues in cells and *in vivo*. In contrast to the general preference for phosphorylation at Ser, some kinases, phosphatases, and protein-interaction domains alternatively have a strong preference for interaction with Thr over Ser.^3-10^ In addition, Thr and Ser phosphorylation sites exhibit different rates of evolutionary change, suggesting functional differences in phosphorylation at each residue.^2,11^

Ser and Thr phosphorylation induce protein-protein interactions that involve specific recognition of proteins with pSer or pThr.^4,5,12-15^ However, pSer and pThr are also capable of directly interacting with the local protein backbone, and thus can directly impact local structure. In their unmodified forms, Ser/Thr free hydroxyls can interact via hydrogen bonding with their own or adjacent amide hydrogens and carbonyls, via their hydroxyl hydrogen bond donor (O–H) and/or acceptors (lone pairs). These interactions lead to significant conformational heterogeneity in Ser and Thr.^16^ In contrast, pSer and pThr only exhibit hydrogen bond acceptors in the dianionic form that dominates at physiological pH (typical p*K*_a_ ∼ 5.5-6.0), suggesting the possibility of changes in backbone interactions due to phosphorylation.

In model peptides, phosphorylation strongly induces α-helix at the N-terminus of the α-helix.^17-19^ In contrast, phosphorylation strongly disrupts α-helix in the interior of the α-helix, including complete disruption of the α-helix in a manner comparable to proline.^19-21^ In proline-rich and intrinsically disordered peptides, circular dichroism (CD) and NMR data indicated substantial conformational restriction in the dianionic phosphorylated residue compared to the unmodified or monoanionic phosphorylated residue.^19,22,23^ Strikingly, these data also demonstrated substantially greater conformational changes due to Thr phosphorylation compared to Ser phosphorylation. Thr, as a β-branched amino acid, generally prefers a relatively more extended conformation.^24,25^ In contrast, dianionic pThr was observed to have a structural signature indicating a highly ordered, compact conformation, including ordered *ϕ* and an intraresidue phosphate-amide hydrogen bond, in these peptides.

Collectively, the data in tau-derived peptides, in proline-rich model peptides, and in α-helical model peptides suggest that pThr induces greater conformational restriction than pSer and induces a greater total structural change as a result of phosphorylation. Given the significance of Ser/Thr phosphorylation in intracellular signaling, and the specific importance of proline-directed phosphorylation in signal transduction, we sought to more broadly examine the structural effects of Ser versus Thr phosphorylation. Of particular note, Ser-Pro and Thr-Pro sequences are phosphorylated by proline-directed kinases, such mitogen-activated protein (MAP) kinases and cyclin-dependent protein kinases (cdks), which are critical in signal transduction pathways and control of cell cycle progression. In addition, some non-proline-directed kinases such as GSK-3β also can phosphorylate at Ser-Pro or Thr-Pro sites. In total, proteomics data indicate that over 25% of Ser/Thr phosphorylated protein sites are pSer-Pro or pThr-Pro sites.^2^ The identification of differences in induced structure for pSer versus pThr at proline-directed phosphorylation sites thus would have broad implications in structural and cellular biology.

## Results

In order to more broadly examine the structural effects of Ser/Thr phosphorylation, a series of peptides derived from proteins important in intracellular signaling and transcription was synthesized (Figure 1). In addition, peptides containing a simple Ser/Thr-Pro dipeptide sequence were examined, in order to identify the inherent effects of phosphorylation on structure in minimal contexts and to determine if the structural changes observed in larger peptides are also found in these minimal proline-directed kinase substrate sequences. These sequences, in combination with those examined previously, represent a wide array of biomedically important proteins containing Ser/Thr-Pro sequences as well as diverse sequence contexts in the residues prior to the phosphorylated residue (acidic, basic, hydrophobic, proline, neutral polar, neutral small nonpolar).

**Figure 1.**
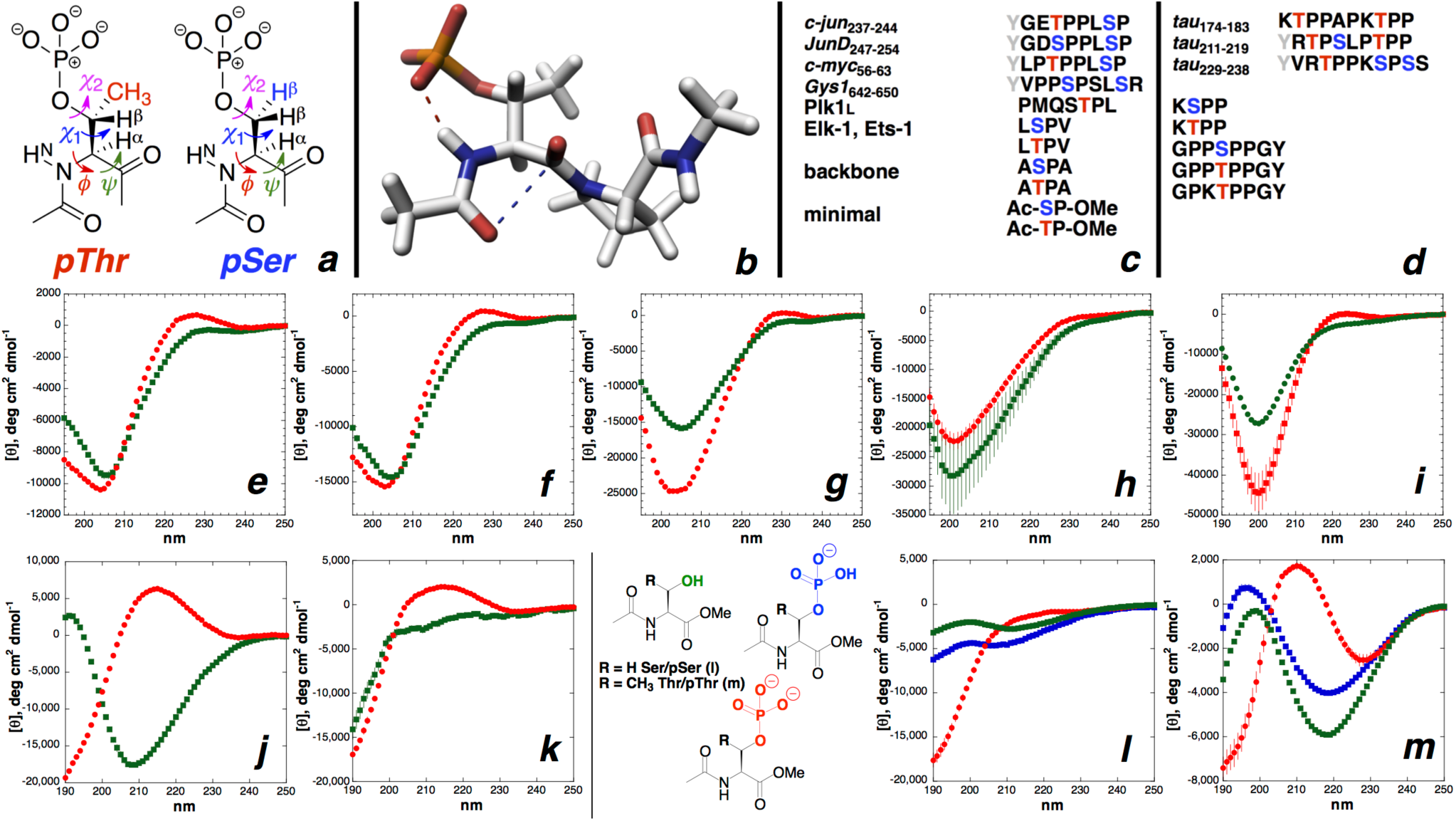
Phosphothreonine (pThr) and phosphoserine (pSer) residues and peptides examined. (a) Definitions of torsion angles and key nuclei analyzed in NMR experiments. Residues adjacent to pSer/pThr are omitted for clarity. (b) Structure of an Ac-pThr-Pro-NHMe dipeptide (from pdb 3a8x,^26^ PKC-ι; pThr *ϕ,ψ* = –51°, +122°, *χ*_1_ = –49° (*g*–), *χ*_2_ = +112°. Hydrogen bond (red dashed line, 2.71 Å O–N distance) and n→π* (blue dashed line, 2.83 Å O–C distance) noncovalent interactions observed in this structure are indicated. Hydrogens were added in Pymol. (c) Peptides examined. Phosphorylated Thr (blue) and Ser (red) residues are indicated. Tyr (grey) were added for concentration determination. All peptides were acetylated on the N-terminus and all peptides except dipeptides contained C-terminal amides. The human proteome (uniprot.org, 20,996 protein sequences (July 20, 2018)) includes 15,893 proteins that contain an SP sequence (68,736 total SP sequences) and 13,927 proteins with a TP sequence (42,271 total TP sequences). 518 and 275 proteins have an ASPA or ATPA sequence; 723 and 441 proteins have a PSPP or PTPP sequence; 374 and 249 proteins have an LSPV or LTPV sequence; 186 and 189 proteins have a KSPP or KTPP sequence; and 205 and 186 proteins have an ESPP or ETPP sequence. (d) Tau-derived and proline-rich peptides examined previously.^23^ (e-i) CD spectra of non-phosphorylated (green squares) and phosphorylated (red circles) peptides. An increase in the magnitude of the positive band around 220 nm indicates an increase in the population of polyproline II helix. (e) c-jun_237-244_; (f) JunD_247-254_; (g) c-myc_56-63_; (h) glycogen synthase-1 Gys1_642-650_; (i) optimized Plk1 polo-box domain ligand (Plk1_L_); (j,k) Neutral (green squares, pH 2) and anionic (red circles, pH 7) (j) Ac-DP-OMe and (m) Ac-EP-OMe; (l,m) Non-phosphorylated (green squares), monoanionic phosphorylated (blue squares, pH 4), and dianionic phosphorylated (red circles, pH 8) (l) Ac-Ser-Pro-OMe and (m) Ac-Thr-Pro-OMe. Experiments were conducted in phosphate-buffered saline at 2 °C (i) or 25 °C (all other peptides).

### Effects of Ser and Thr phosphorylation by CD spectroscopy

All peptides were examined by CD in the non-phosphorylated and phosphorylated forms. Across all peptides containing pSer-Pro or pThr-Pro sequences, phosphorylation was observed by CD to result in an increase in polyproline II helix (PPII), which is indicated by an increase in the magnitude of the positive band at ∼228 nm (Figure 1e-i).^27-29^

The effects of phosphorylation were examined in Ac-Ser-Pro-OMe and Ac-Thr-Pro-OMe dipeptides (Figure 1lm). The effect of “pseudophosphorylation” (mutation of a phosphorylatable Ser/Thr to Asp or Glu) was also examined, using the peptides Ac-Asp-Pro-OMe and Ac-Glu-Pro-OMe.^30,31^ The CD spectrum of the non-phosphorylated Thr-containing peptide exhibited an intense negative band *λ*_min_ = 219 nm, consistent with extended and/or turn conformations that is expected for the β-branched residue Thr. The CD spectrum with Ser was less well-defined and consistent with greater disorder. In contrast, both phosphorylated peptides when in the dianionic state exhibited an increased PPII structural signature (more positive signal at 210-220 nm)^28^ and a significant change in structure from the nonphosphorylated peptides, indicating that induction of order is an inherent property of phosphorylation at Ser/Thr-Pro sequences. Notably, the monoanionic phosphorylated peptides exhibited only modest changes compared to non-phosphorylated Ser or Thr, indicating that the structural effects of phosphorylation are primarily mediated via the dianionic ionization state.^23^

Across these peptides, Thr phosphorylation both induced a more robust PPII structure and a more dramatic total change in structure upon phosphorylation, with stronger CD signatures in both the non-phosphorylated and phosphorylated states. These data indicate a greater disorder-to-order change in structure upon Thr phosphorylation than upon Ser phosphorylation, and suggest Thr phosphorylation sites as having greater importance than Ser phosphorylation sites for induced structural effects of phosphorylation.

### Effects of Ser and Thr phosphorylation by NMR spectroscopy

To identify residue-specific structural changes upon phosphorylation, all peptides were examined by ^1^H and ^15^N NMR spectroscopy, comparing the effects of phosphorylation on Ser versus Thr residues on chemical shift (δ) and on the *ϕ* backbone torsion angle (via ^3^*J*_αN_, the coupling constant between H^N^ and Hα, and a parametrized Karplus relationship^32^). For a subset of peptides, additional analysis was conducted via ^1^H-^13^C HSQC experiments, which provide residue-specific ^13^C chemical shifts, which are more sensitive to conformation and less sensitive to local electrostatic environment than ^1^H chemical shifts.^33^ In addition to peptides analyzed above by CD, peptides were also examined to compare Ser versus Thr phosphorylation in alternative local sequence contexts, including a “backbone-only” context lacking side chains beyond the β carbon on adjacent residues (Ac-A(S/T)PA-NH_2_) and a context with hydrophobic residues adjacent to the phosphorylation site (Ac-L(S/T)PV-NH_2_; these sequences are found as phosphorylated sites in the transcription factor Elk-1, the proto-oncogene transcription factor Ets-1, the RNA-splicing regulatory protein AHNAK (12 LSPV phosphorylation sites), and numerous other proteins).

Across all peptides, phosphorylation resulted in a large downfield change in the δ of the pSer/pThr amide hydrogens (H^N^) and amide nitrogens (N^H^), as well as a reduction in the ^3^*J*_α_N coupling constants (Figure 2, Figure 3, Table S1). These data indicate a significant increase in order and the adoption of a more compact conformation at the *ϕ* torsion angle upon phosphorylation (Thr average *ϕ* = –84°, pThr average *ϕ* = –55°; Ser average *ϕ* = –80°, pSer average *ϕ* = –71°). ^3^*J*_αN_ values between 6-8 Hz are typical for disorder and can result from significant conformational averaging, whereas ^3^*J*_αN_ < 6 Hz have unique solutions when –165° < *ϕ* < –30° (i.e. in the allowed region of the left half of the Ramachandran plot), and thus require a compact *ϕ* as the major conformation, with smaller values indicating greater order. The downfield amide chemical shifts, as well as the observation of the amide hydrogens at pH 7.5-8.0 (a pH at which amide hydrogens in disordered proteins usually exchange rapidly and therefore are not observed by NMR), indicate amide protection and suggest a close interaction with the phosphate via a hydrogen bond.^23^ Further evidence of a hydrogen bond is indicated by the small pThr H^N^ temperature coefficient (Δδ < 5 ppb).^34^

**Figure 2.**
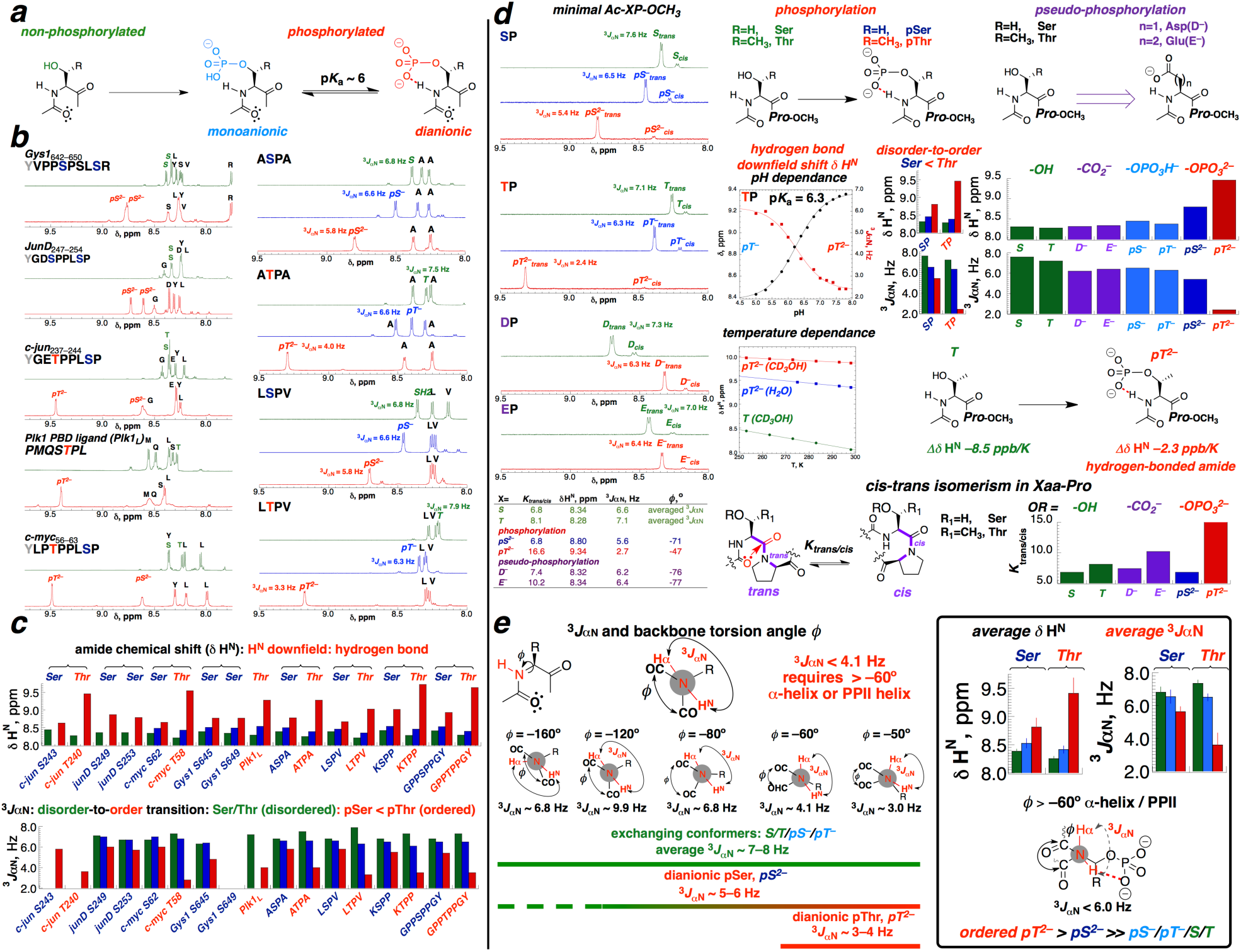
^1^H NMR data on peptides as a function of phosphorylation and ionization state. (a) Phosphorylation and ionization states of Ser and Thr. (b) 1-D NMR spectra of peptides. (c) Comparison of amide H chemical shifts (H^N^ δ) and H^N^-Hα coupling constants (^3^*J*_αN_). (d) NMR data on minimal XP dipeptides. (e) Summary of average H^N^ δ and ^3^*J*_αN_.

**Figure 3.**
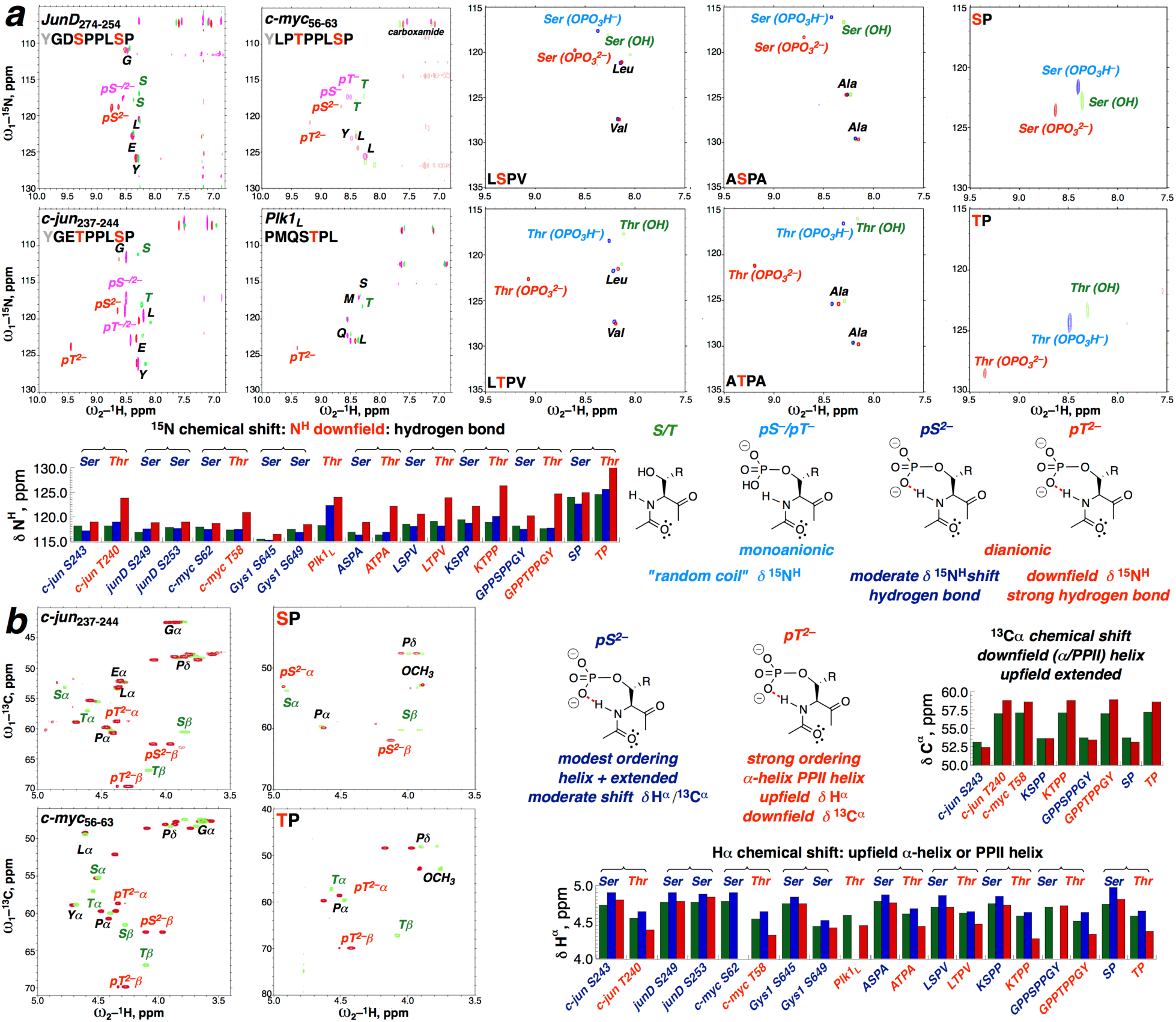
NMR data on peptides from 2-D heteronuclear experiments. (a) ^1^H-^15^N HSQC spectra and data on amide hydrogen (H^N^) and amide nitrogen (N^H^) chemical shifts. (b) ^1^H-^13^C HSQC spectra and Hα and Cα chemical shifts.

Notably, the structural signatures observed by NMR in diverse protein-derived peptides were similarly observed in the minimal pSer-Pro and pThr-Pro dipeptides, suggesting that minimal peptides provide a general insight into the inherent effects of proline-directed phosphorylation. At all Ser/Thr-Pro sites, the induced structures at pThr were more defined, more compact, and more ordered than the induced structures at pSer, as indicated by more downfield H^N^ and N^H^ δ, smaller ^3^*J*_αN_ coupling constants (indicating a more compact and more ordered structure)^32^, and larger changes in Hα and Cα δ (which correlate to main chain conformation)^33,35,36^ for pThr residues. Moreover, comparison of the closely related c-jun (with one pThr and one pSer) and JunD (two pSer) peptides, as well as the Ac-pSer-P-OMe versus Ac-pThr-P-OMe peptides, and the A(S/T)PA and L(S/T)PV tetrapeptides, emphasizes the clear differentiation between the effects of pSer and pThr on amide ^1^H and ^15^N chemical shifts.

The magnitude of the structural effects of phosphorylation was strongly dependent on the dianionic form of pThr: monoanionic pThr and monoanionic pSer (data at pH 4) had only modest structural effects compared to the unmodified peptides, including small downfield changes in amide δ and small decreases in ^3^*J*_αN_. Indeed, ^3^*J*_αN_ for monoanionic pSer and pThr are closer to those of the pseudophosphorylation mimics Asp or Glu in model peptides, where only modestly induced order was observed. Combining the more extended conformations observed in general at β-branched Thr residues compared to Ser residues with the significantly more compact structures observed herein at pThr residues, these data collectively indicate that larger overall induced structural changes are observed on Thr phosphorylation (extended to compact conformation) than on Ser phosphorylation. Moreover, these data suggest that pseudophosphorylation can only partially replicate the structural effects of phosphorylation, and that pseudophosphorylation more effectively mimics pSer than pThr.

We previously observed that the activation loop pThr197 of protein kinase A (PKA) (pdb 1rdq,^37^ 1.26 Å, local sequence RTW**pT**LCG) exhibited a structure similar to that found in proline-rich peptides, including PPII *ϕ,ψ* torsion angles, a hydrogen bond to its own amide hydrogen, a *g*^−^ conformation in *χ*_1_, and, surprisingly, an eclipsed C–Hβ/Oγ–P bond (*χ*_2_ ∼ +120°). The CD and NMR spectra of pThr and, to a lesser extent, pSer in peptides herein were consistent with the main chain conformation observed crystallographically in PKA. To identify the side chain conformational changes upon phosphorylation, selected peptides were analyzed by NMR (Figure 4) to determine the coupling constant ^3^*J*_αβ_, which reports on the *χ*_1_ torsion angle, whose maximum value of 10 Hz indicates an *anti*-periplanar relationship of the hydrogens and thus a *g*^−^ conformation via a parametrized Karplus relationship;^38,39^ and ^3^*J*_PHβ_, which correlates to the *χ*_2_ torsion angle (local maximum of 10.5 Hz at χ_2_ = +120° with an eclipsed/*syn*-periplanar relationship of the Oγ–P and C–Hβ bonds).^40^ Across all peptides examined, the ^3^*J*_αβ_ coupling constants increased upon Thr phosphorylation (Thr mean ^3^*J*_αβ_ = 6.4 Hz, pThr mean ^3^*J*_αβ_ = 9.2 Hz, data from 4 peptides). In pThr residues, both ^3^*J*_αβ_ and ^3^*J*_PHβ_ (mean pThr ^3^*J*_PHβ_ = 8.8 Hz) approached the maxima expected if the crystallographically observed conformations were present in the peptides, a surprising result given the inherent disorder in these sequences, which are not stabilized by tertiary structure. In contrast, the ^3^*J*_PHβ_ values for the diastereotopic pSer β hydrogens were smaller (mean pSer ^3^*J*_PHβ_ = 5.9 Hz) and not distinct in any condition examined, consistent with greater *χ*_2_ conformational hetereogeneity in pSer residues.

**Figure 4.**
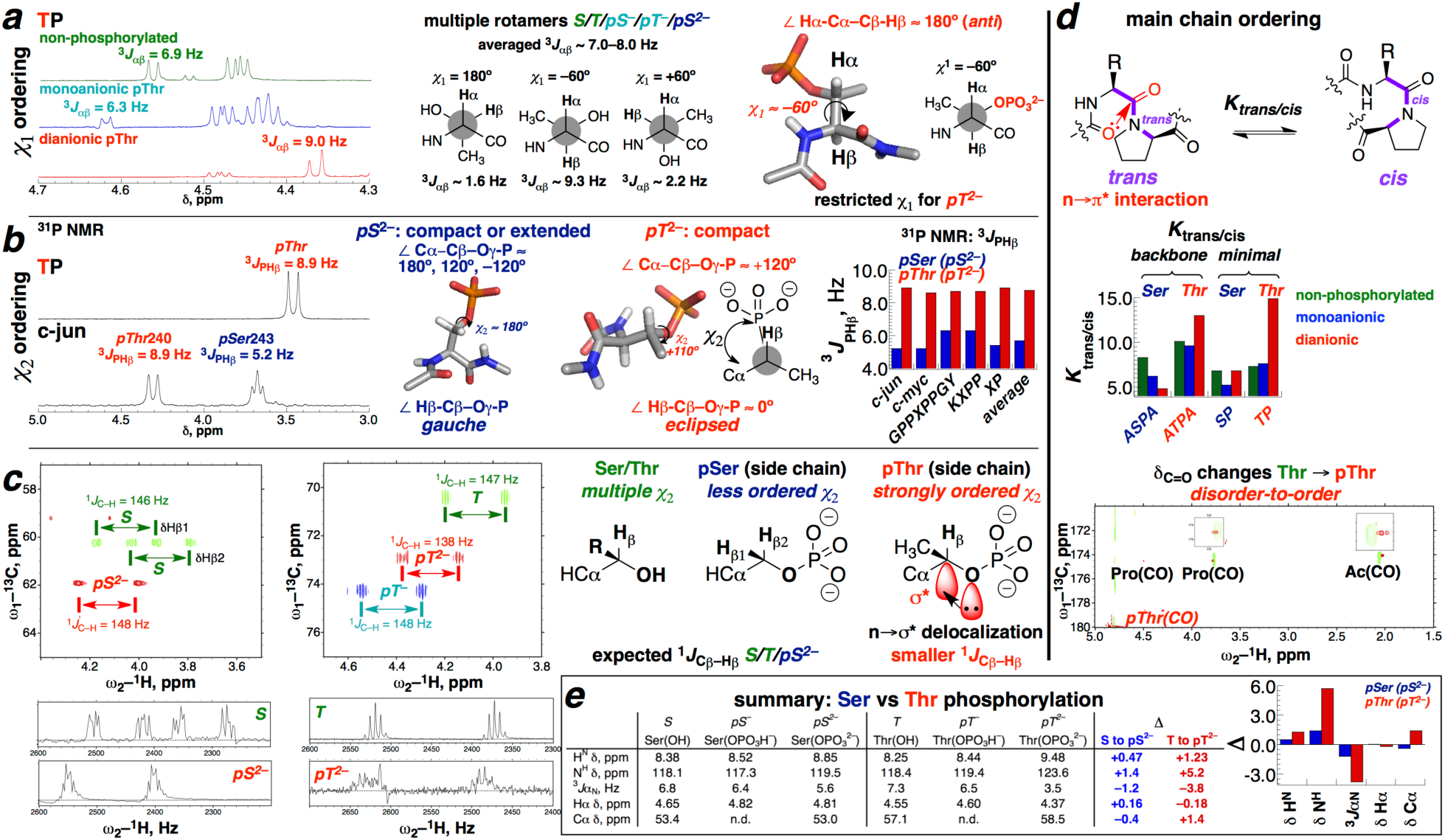
NMR data to identify the effects of phosphorylation on torsion angles of the side chain (*χ*_1_ and *χ*_2_) and main chain (*ω*). (a) ^1^H NMR spectra (Hα region) in 100% D_2_O, with ^3^*J*_αβ_ indicated. (b) ^31^P NMR spectra in 100% D_2_O, with ^3^*J*_PHβ_ quantified. (c) Effects of phosphorylation on 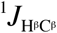. (d) Effects of phosphorylation in model peptides on *trans* versus *cis* proline amide bond (*K*_trans/cis_; *ω* torsion angle) and on carbonyl ^13^C δ. (e) Summary of NMR data across all peptides in Figure 1cd as a function of amino acid (Ser vs Thr) and phosphorylation and ionization states.

### Analysis of pThr and pSer in the Protein Data Bank (PDB)

The results above, combined with prior work, suggest that structural changes induced by phosphorylation on Ser and Thr are general, with a particularly strong conformational bias induced by dianionic pThr. To examine this generality, pSer and pThr residues in the PDB were examined, and the conformational preferences of these phosphorylated residues compared to non-phosphorylated Ser and Thr from high-resolution structures (≤ 1.25 Å resolution, ≤ 25% sequence identity). Due to a limited number of pSer and pThr residues in the PDB, which is a caveat to the PDB analysis that follows, phosphorylated proteins were examined with expanded resolution (≤ 2.50 Å) and sequence identity (95%) cutoffs, yielding 234 pSer and 155 pThr residues that were analyzed, including residues in globular proteins and ligands in protein-protein complexes (Tables S2, S3).

These data indicate a remarkable degree of order at pThr residues, compared to non-phosphorylated Thr or Ser, or to pSer (Figure 5). The Ramachandran plot of pThr exhibits a strong preference for highly ordered values of *ϕ*, with PPII the major conformation and α-helix a minor conformation. These data are consistent with the conformational preferences observed herein and previously in diverse sequence contexts, where pThr exhibited a small ^3^*J*_αN_ indicative of a high degree of order in *ϕ* and consistent with either PPII or α-helix. In contrast, Thr in general adopts a more extended and/or more disordered conformation due to β-branching of the sidechain and the possibility of multiple sidechain-main chain hydrogen bonds. pSer residues in the PDB exhibit greater order than Ser, with similar but less distinct preferences for PPII and α-helix, but weaker conformational preferences than pThr. The conformational effects of Thr versus Ser phosphorylation were particularly notable when examining the *ϕ, χ*_1_, and *χ*_2_ torsion angles individually. pThr exhibits a strong preference for *ϕ* ∼ –60°, in contrast to the more extended *ϕ* preference of Thr and the more diffuse *ϕ* preference of pSer. pThr also exhibits a strong *χ*_1_ preference for the *g*^−^ conformation, in contrast to the similar *g*^−^ and *g*^+^ preferences of Thr and pSer, and the absence of a significant *χ*_1_ preference for Ser.^16^

**Figure 5.**
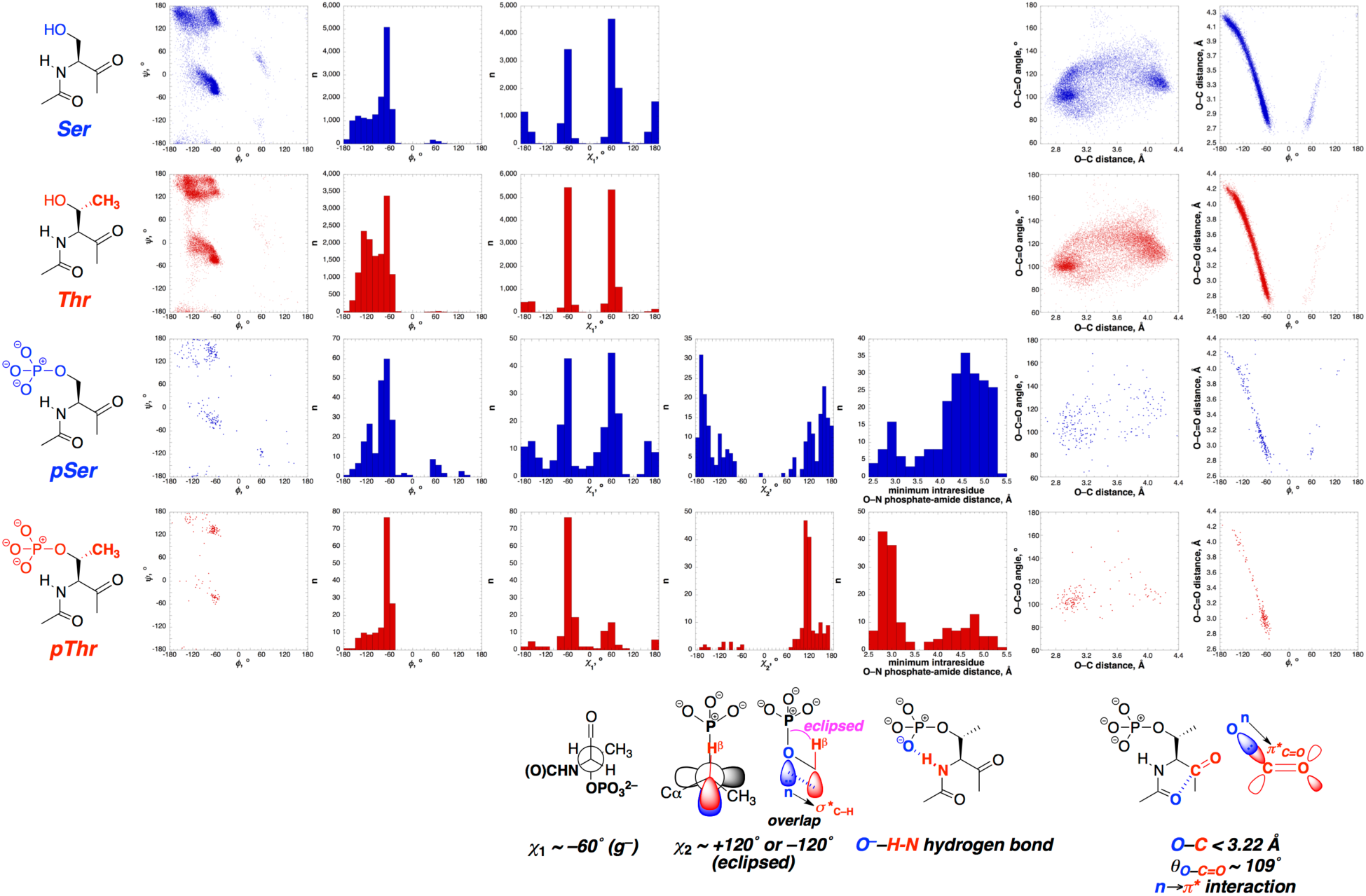
Analysis of Ser (top, blue), Thr (top, red), pSer (bottom, blue), and pThr (bottom, red) residues in the PDB. From left to right: amino acid structure; Ramachandran plot; histograms of *ϕ, χ*_1_, and *χ*_2_ torsion angles; histogram of shortest distance between any phosphate oxygen and the amide nitrogen of the same residue; n→π* parameters O_*i*_–C_*i*+1_ angle versus O_*i*_–C_*i*+1_ distance between consecutive carbonyls; O_*i*_–C_*i*+1_ intercarbonyl n→π* distance versus *ϕ* main chain torsion angle.

Similarly, pThr strongly prefers *χ*_2_ ∼ +120°, with eclipsing or near-eclipsing Oγ–P and C– Hβ bonds. Interestingly, eclipsing Oγ–P and C–Hβ bonds are also observed (*χ*_2_ ∼ +120° or – 120°) as minor conformations of pSer, indicating that the eclipsed conformation is energetically favorable, although for pSer the sterically preferred fully extended *χ*_2_ ∼ 180° is the major conformation. The low population of conformations with negative values of *χ*_2_ is not surprising for pThr, as these conformations would induce significant steric repulsion of the large phosphate group with the γ-CH_3_ group and/or the protein main chain. However, the apparent preference for an eclipsed conformation was not expected for pThr, although a similar interaction has been observed by NMR in glycosylated Thr.^41,42^ The greater length of the O–P bond (1.62 Å) compared to the C–H bond (1.09 Å) is expected to reduce, but not eliminate, the steric cost of an eclipsing interaction (H…P distances 2.5–2.6 Å, less than the 3.0 Å sum of the van der Waals radii of P and H).

Further analysis indicates a Cβ–Oγ–P bond angle of ∼120° in these structures (i.e. sp^2^-type geometry at Oγ), indicating that the sp^2^-like O lone pair and the Cβ-Hβ bond lie within the same plane, with the Oγ sp^2^-like lone pair and the σ* of C–Hβ aligned for potential orbital overlap. The unusual conformational preference for an eclipsed conformation of *χ*_2_ is herein ascribed to the presence of a favorable n→σ* hyperconjugative interaction between the coplanar sp^2^-like Oγ lone pair (n) of the electron-rich dianionic pThr and the antibonding orbital (σ*) of the electron-deficient C–Hβ bond. This type of n→σ* interaction (known as the Perlin effect) is observed at H–C–O–: torsion angles in carbohydrates when the C–H is axial and thus *anti*-periplanar to the oxygen lone pair (i.e. the oxygen lone pair is *syn*-periplanar with the C–H σ*, as is observed herein).^43-45^ In these stereoelectronic effects, the 1-bond C–H coupling constant ^1^*J*_CH_ is observed to be 8-10 Hz smaller than in similar C–H bonds that are not stabilized by this interaction. Indeed, consistent with this interpretation, 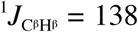 Hz for pThr in dianionic Ac-pTP-OMe, compared to 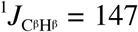 Hz in non-phosphorylated Ac-TP-OMe and 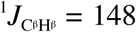 Hz in monoanionic Ac-pTP-OMe (Figure 4), similar to a typical Perlin effect and indicating a significant interaction only when pThr is in the more electron-rich dianionic state, where the O lone pair can function as an effective lone pair donor. These results suggest that stabilization of the *χ*_2_ eclipsed conformation occurs via an n→σ* stereoelectronic effect.

pThr exhibited a particularly strong preference for a hydrogen bond between the phosphate and the pThr amide hydrogen. In contrast, while pSer also exhibited a phosphate-amide hydrogen bond as a significant preference, pSer exhibited a substantially lower preference for an intraresidue phosphate-amide hydrogen bond, consistent with its preference for the sterically preferred fully extended *χ*_2_. The PDB data are consistent with the observations herein of more downfield H^N^ and N^H^ amide δ and slower amide exchange at pH 7.5-8.0 (as evidenced by less broadening) for pThr amides compared to pSer amides in peptides.

n→π* interactions between adjacent carbonyls have recently emerged as a significant force stabilizing local interactions between consecutive residues.^46-52^ n→π* interactions involve electron delocalization via orbital overlap between the lone pair of a donor carbonyl oxygen (n refers to the lone pair, on the *i* residue) and the antibonding (π*) orbital of the subsequent carbonyl (*i*+1 residue). An n→π* interaction can be identified by the presence of an O_*i*_–C_*i*+1_ interresidue distance that is less than the sum of the C and O van der Waals radii (d < 3.22 Å, with shorter distances indicating greater orbital overlap) and an O_*i*_–C_*i*+1_=O angle ∼109°, that maximizes orbital overlap. Data from the PDB indicate a strong preference for pThr residues to exhibit n→π* interactions between the carbonyl prior to pThr (i.e. the carbonyl directly conjugated to the pThr amide N–H) and the pThr carbonyl (Figure 5). The net effect of this interaction is to induce strong local ordering of two consecutive carbonyls.

Analysis of pSer residues indicated a relatively reduced likelihood of n→π* interactions and less clustering around the ideal 109° angle for n→π* interactions. Notably, across all residues, there was a strong correlation of O–C interresidue bond length with the *ϕ* torsion angle, consistent with n→π* interactions being more favorable in more compact conformations.^48,49^ NMR data on Ac-pTP-OMe and KpTPP peptides were also consistent with an n→π* interaction stabilizing the structure of these peptides, with significant changes in the acetyl and pThr carbonyl ^13^C δ in the phosphorylated peptides over the non-phosphorylated peptides (Figure 4c). Interestingly, in the minimal Ac-pTP-OMe peptide, as well as in other tetrapeptides with TP sequences, phosphorylation also induced a significant increase in preference for *trans* over *cis* amide bonds when pThr was dianionic (Figure 4d). In contrast, in Ac-pSP-OMe and other peptides with dianionic pSer, an increase in the population of *cis*-proline amide conformation was observed compared to the unmodified Ser. These data suggest an inherent difference in the propensity of pSer and pThr to promote a *cis* amide bond.^53^ The increased preference for a *trans* amide bond with pThr could potentially be due to the enhanced n→π* interaction in the phosphorylated peptide, whereby the intraresidue hydrogen bond makes the conjugated carbonyl more electron-rich (more δ^−^), and therefore a better donor for an n→π* interaction.

### X-ray crystallography of a phosphothreonine-proline dipeptide

The structural signatures observed for pThr in the PDB were consistent with NMR data observed in minimal Ac-pTP-OMe dipeptides. To further examine the structural effects of Thr phosphorylation, a derivative of this peptide was synthesized with an iodobenzoyl acyl group (4-I-Bz-pThr-Pro-NH_2_) and a C-terminal amide. In addition, this peptide was independently synthesized in both the _L_-peptide and D-peptide enantiomeric forms, to allow for an enhanced possibility of crystallization via racemic crystallization.^54^

Both the single-enantiomer (l-peptide) and the racemic (1:1 mixture of l-peptide and d-peptide) solutions of 4-I-Bz-pThr-Pro-NH_2_ crystallized via vapor diffusion. The single-enantiomer crystal structure was solved at 0.80 Å resolution (Figure 6). The racemic crystal structure was solved at 0.87 Å resolution, with two non-equivalent molecules in the asymmetric unit. The crystal structures recapitulated all of the structural features identified via NMR spectroscopy across diverse peptide sequences and via bioinformatics analysis. In all three independent molecules across the two crystal structures, pThr adopted a PPII conformation with a compact value of *ϕ*, a close intraresidue phosphate-amide hydrogen bond, and a close intercarbonyl O_ac_…C_pThr_=O distance, consistent with a favorable n→π* interaction stabilizing this conformation. In all structures, the side chain exhibited a *g*– *χ*_1_ conformation and an eclipsed *χ*_2_ conformation. Interestingly, in the single-enantiomer structure, a water molecule was localized by hydrogen bonding between the amide-interacting phosphate oxygen and the pThr carbonyl. Hydrogen bonding at the acceptor carbonyl would be expected to enhance the interaction strength of an n→π* interaction. The bound water observed herein could potentially be an additional factor by which solvation associated with Thr phosphorylation stabilizes the PPII conformation.

**Figure 6.**
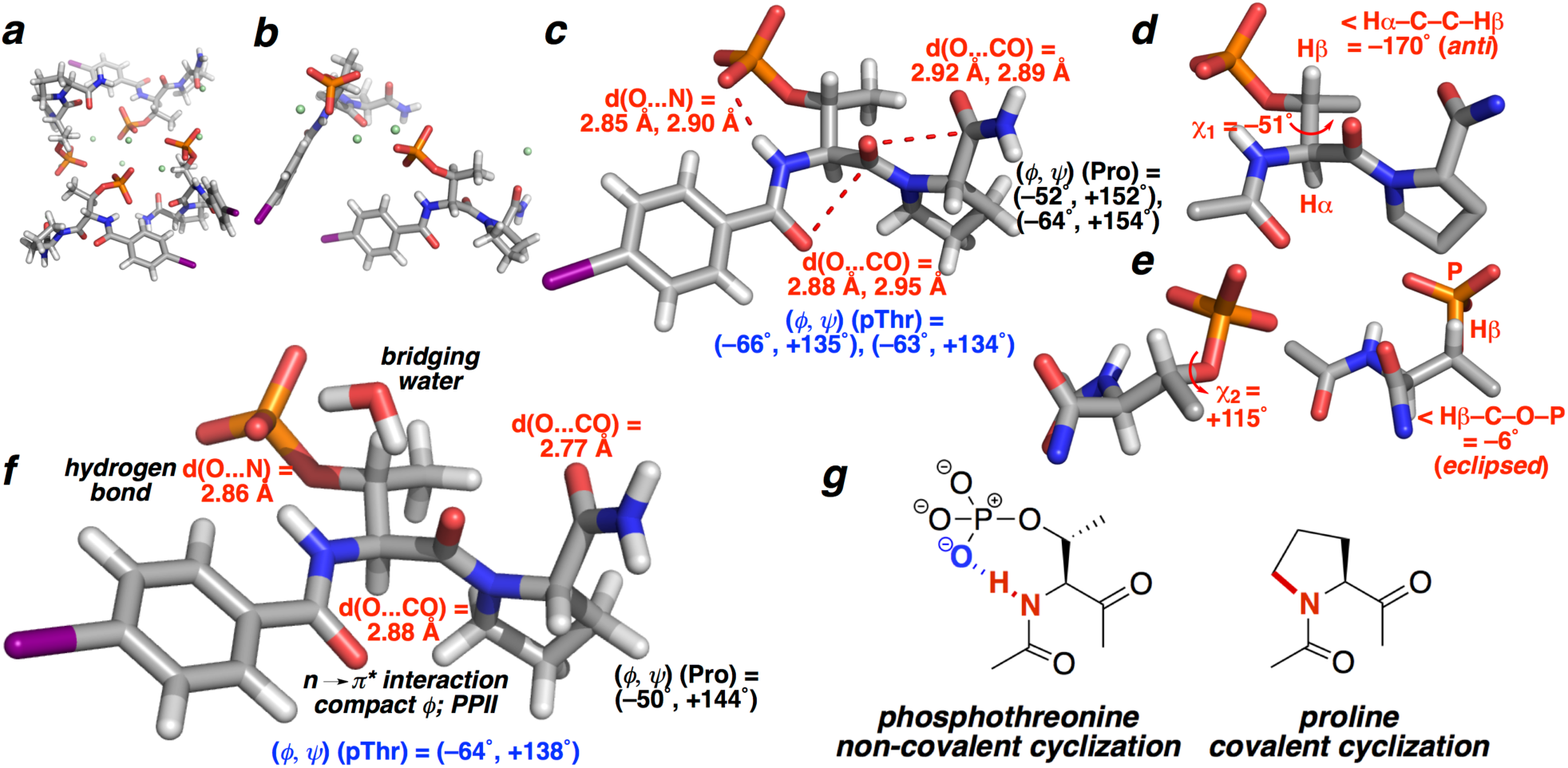
Structures of dianionic (a-e) (±)-4-I-Bz-pThr-Pro-NH_2_ (resolution 0.87 Å) and (f) single-enantiomer 4-I-Bz-pThr-Pro-NH_2_ (resolution 0.80 Å), as determined by X-ray crystallography. (a) Crystal packing showing 4 molecules, two l-peptides and two d-peptides (the asymmetric unit plus its mirror image). Water molecules (10 per asymmetric unit) found in the crystal structure are omitted for clarity. Sodium atoms are indicated in green. (b) Asymmetric unit (P_–1_ centrosymmetric space group), containing one molecule of l-peptide and one of d-peptide, with distinct (not symmetrically related) structures. (c) Structure of one molecule of 4-I-Bz-pThr-Pro-NH_2_, with torsion angles and distances from both structures indicated. (d,e) Analysis of (d) *χ*_1_ and (e) *χ*_2_ torsion angles in one molecule of the structure. The torsion angles of the _D_-peptide were inverted to those of its _L_-peptide equivalent for clarity. (f) Single-enantiomer crystal structure of 4-I-Bz-pThr-Pro-NH_2_. Additional water molecules and sodium and potassium ions are omitted for clarity. In addition, the structures indicate significant puckering at the pThr and Pro carbonyls, but not at the iodobenzoyl carbonyl, consistent with the participation of the acceptor carbonyls in an n→π* interaction. The bridging water molecule observed in (f) exhibits close (2.9 Å O…O distances) hydrogen bonds with both the phosphate oxygen and the pThr carbonyl. To our knowledge, this bridging water molecule has not previously been observed. (g) Comparison of the non-covalently cyclic preferred structure of pThr and the covalently cyclic structure of Pro.

The strong intraresidue hydrogen bond for dianionic pThr observed crystallographically results in a non-covalently cyclic structure of pThr, analogous to the covalently cyclic structure of proline (Figure 6g). Consistent with this interpretation, we previously observed that dianionic pThr is strongly destabilizing to α-helices when located at the center and C-terminus of the α-helical sequence, in a manner comparable to the effects of Pro on α-helices.^19,20^ The crystallographic data also indicate an n→π* interaction between the iodobenzoyl carbonyl and the pThr carbonyl. An n→π* interaction is further observed between the pThr carbonyl and the Pro carbonyl, with the Pro exhibiting an *exo* ring pucker and PPII conformation. These data indicate the ability of Thr phosphorylation to organize multiple consecutive residues. Notably, these non-covalent interactions are expected to be synergistic in leading to an ordered structure: a phosphate-amide hydrogen bond would be expected to increase electron density on the conjugated carbonyl, making it a better donor for an n→π* interaction with the subsequent carbonyl.^51^

These data represent the first small-molecule X-ray crystallographic data on a peptide containing pThr. The X-ray structure of a simple pThr-containing peptide indicates that dianionic pThr has strong inherent conformational preferences that are manifested in a side chain-main chain intraresidue hydrogen bond, restrained *ϕ, ψ, χ*_1_, and *χ*_2_ torsion angles, and a favorable interresidue n→π* interaction that organizes consecutive residues. Moreover, these data suggest that the similar conformational preferences observed for pThr residues in the PDB, in diverse sequence contexts, reflect the strong inherent residue preferences of dianionic pThr.

### Computational investigations on minimal peptides containing pThr

To further characterize the molecular basis of the highly stabilized structure of pThr within peptide and protein contexts, geometry optimization calculations were conducted by density functional theory (DFT) methods on a minimum Ac-pThr-NMe_2_ structure, on pThr-Pro dipeptides, on variants of these with the bridging water molecule observed in the single-enantiomer crystal structure, and on the equivalent peptides with pSer in place of pThr.

Geometry optimization calculations yielded structures (Figure 7, Table S4) that were similar to those observed crystallographically and to the conformational features observed in solution by NMR spectroscopy. In particular, across all peptides examined, the resultant structures exhibited *ϕ* and *ψ* torsion angles and other conformational features that were very similar to those observed crystallographically. In addition, the data indicated significant pyramidalization at the pThr carbonyl (2.2–3.0°), but not at the acetyl carbonyl (≤ 0.3°). This pyramidalization is consistent with an important role of the n→π* interaction in pThr stabilizing the PPII conformation, whereby the acceptor carbonyl exhibits partial pyramidalization analogous to the early stages of a nucleophilic attack of one carbonyl on the other.^46,47,49,52^ Interestingly, equivalent geometry-optimized structures with pSer exhibited somewhat more extended conformations and longer n→π* interaction distances even with the same overall conformational features, consistent with the reduced preference for a compact conformation for pSer compared to pThr.

**Figure 7.**
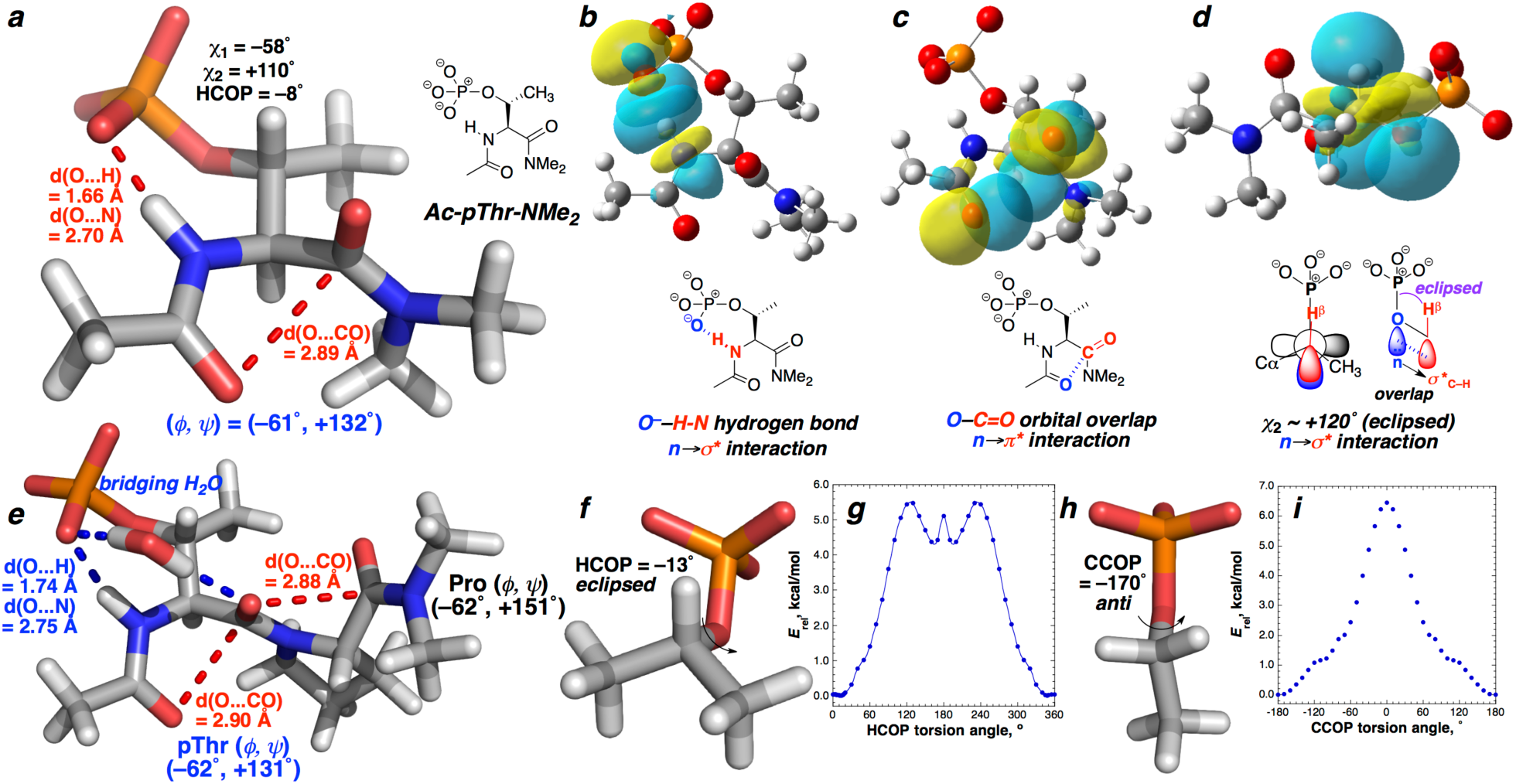
(a) Structure resulting from geometry optimization of Ac-pThr-NMe_2_ using the M06-2X method and the 6-311++G(3d,3p) basis set in implicit H_2_O. (b-d) Natural bond orbital (NBO) analysis of Ac-pThr-NMe_2_, indicating the molecular orbital basis of stabilizing non-covalent interactions in pThr. Stabilization is provided by overlap of orbital lobes. Favorable electrostatics also contribute to stabilization. Via NBO analysis, the n→π* interaction is stabilized by 1.1 kcal mol^−1^ via electron delocalization between the acyl carbonyl p-like lone pair and the pThr carbonyl π* molecular orbitals, while the n→σ* interaction that stabilizes the eclipsed conformation is favorable by 4.7 kcal mol^−1^. (e) Structure resulting from geometry optimization of Ac-pThr-Pro-NMe_2_•H_2_O. (f-i) Geometry-optimized structures (f, h) and torsion angle scans equivalent to *χ*_2_ (g,i) on dianionic (f, g) isopropyl phosphate (pThr model) and (h, i) ethyl phosphate (pSer model) using the MP2 method and (f,h) 6-311++G(3d,3p) or (g,i) 6-311++G(2d,2p) basis sets in implicit H_2_O.

Natural bond orbital (NBO)^55^ analysis was conducted on the minimal Ac-pThr-Nme_2_ structure (Figure 7b-d). Structural analysis revealed strong orbital overlap between the nonbonding phosphate oxygen lone pair and the amide N–H σ* in the phosphate-amide hydrogen bond, as expected. Similarly, the donor *i*–1 carbonyl and the acceptor pThr carbonyl exhibited strong orbital overlap between the O_*i*–1_ lone pair (n) and π* of the carbonyl (Figure 7c), as previously seen for favorable n→π* interactions.^49,56^

In addition, significant stabilization energy and orbital overlap were observed between the Oγ lone pair (n) and the antibonding orbital (σ*) of the C–Hβ bond (Figure 7d), indicating that a favorable n→σ* interaction is associated with the preference for an eclipsed conformation at *χ*_2_. Analysis of the energetics of bond rotation about the Cβ-Oγ bond, using a minimal isopropyl phosphate model (Figure 7fg), indicated that, despite unfavorable sterics, the eclipsed conformation (*χ*_2_ ∼ +120°) is the minimum energy conformation at the *χ*_2_ torsion angle, consistent with this conformation being the dominant *χ*_2_ for pThr. In addition, eclipsed conformations (*χ*_2_ = +120° or –120°) were also observed to be favorable minor conformations for an ethyl phosphate model of pSer (Figure 7hi), consistent with bioinformatics data. In sum, these calculations indicate, consistent with experimental data, that dianionic pThr is a particularly ordered amino acid.

## Discussion

We have conducted a comprehensive analysis of the structural effects of Ser and Thr phosphorylation, using a combination of solution analysis of peptides by CD and NMR spectroscopies, bioinformatics analysis of pSer and pThr residues in the PDB, small molecule X-ray crystallography of a pThr-Pro dipeptide, and calculations on minimal peptides containing pThr. Across all size scales and by all methods employed, dianionic pThr was observed to be highly ordered, exhibiting a strong preference for a compact *ϕ* torsion angle (*ϕ* ∼ –60°). This degree of ordering in *ϕ* is unique among all examined canonical and post-translationally modified amino acids, suggesting that dianionic pThr has defined structural roles. The ordering in dianionic pThr is supported by three non-covalent interactions, including a strong intraresidue phosphate-amide hydrogen bond, an n→π* interaction between the carbonyls of adjacent residues that results in ordering of both residues, and an n→σ* interaction between the phosphate γ-O lone pair (n) and the σ* of the C–Hβ that restricts the conformation of *χ*_2_. The overall effect of this combination of interactions is a non-covalently bicyclic structure that fundamentally restricts dianionic pThr to preferentially adopt either a PPII or α-helix conformation. Notably, the non-covalently cyclic structure of dianionic pThr is analogous to the covalently cyclic structure of Pro (Figure 6g), consistent with the similar position-dependent effects of dianionic pThr and Pro on α-helix stability and in inducing PPII.^19,20,22,23^

The structural effects observed herein are dependent on the dianionic state of pThr that predominates at physiological pH. Only modest structural changes were observed for monoanionic pThr. Similarly, the (monoanionic) pseudophosphorylation mimics Asp and Glu were not capable of replicating all of the structural effects of dianionic pThr, exhibiting structures that were intermediate between unmodified Thr and dianionic pThr. In addition to electrostatic differences, these data provide a fundamental structural basis for the inability of pseudophophosphorylation mimics to fully replicate the functional effects of phosphorylation in all cases, particularly for phosphorylation at Thr.

In contrast, the structural effects of phosphorylation at Ser were relatively more modest. While dianionic pSer can adopt a structure with all of the features of dianionic pThr, including an eclipsed C–H/O–P bond (*χ*_2_ ∼ +120° or –120°), the absence of a γ-methyl group results in greater inherent conformational heterogeneity, including a much wider range of conformations at *ϕ, ψ, χ*_1_, and *χ*_2_ and a substantially reduced likelihood of an intraresidue phosphate-amide hydrogen bond. In addition, as was observed with pThr, the structural effects of pSer were more modest in the monoanionic state. Moreover, the non-phosphorylated states of Ser and Thr exhibit substantially different conformational preferences, due to multiple possible side chain-main chain hydrogen bonds and the general preference for a more extended conformation for the β-branched amino acid Thr.^16,24,25^ Consistent with previous analyses, these differing inherent conformational preferences of Ser and Thr are observable by NMR, by CD, and within the PDB.

Moreover, the greater conformational restriction at dianionic pThr compared to pSer or to either Asp or Glu suggests that pseudophosphorylation is more likely to be effective at Ser than Thr phosphorylation sites. Although pseudophosphorylation partially or fully rescues function at Ser and Thr sites in numerous systems, it is also ineffective in many cases. The effects of phosphorylation versus pseudophosphorylation have been examined within protein kinase A (PKA).^57^ Notably, Thr is exceptionally conserved in PKA orthologs that are widely divergent evolutionarily, despite the pThr γ-methyl group being solvent-exposed in crystal structures of activated PKA. In protein kinases with active site phosphorylation, typically the activation loop is disordered, with no electron density, in the non-phosphorylated state, whereas phosphorylation induces order, resulting in electron density for the entire activation loop.^58^ These effects have generally been ascribed to electrostatic and hydrogen bonding interactions with Arg or Lys residues. We propose that the inherent ordering at the phosphorylation site is central to the conformational changes in active site phosphorylation, providing a structurally rigid nucleus for other longer-range interactions that further stabilize the structure of the activation loop. Moreover, we hypothesize that the inherent ordering is greater in protein kinases containing activation loop Thr phosphorylation than in activation loop Ser phosphorylation. Notably, among the human protein kinases, Thr is more common that Ser at known or predicted activation loop phosphorylation residues, despite the inherently greater frequency of Ser phosphorylation.

Ser/Thr phosphorylation and multiple-site phosphorylation functional effects have been examined in numerous contexts. For example, phosphorylation of the transcription factor p53 at multiple Ser sites leads to a graded enhancement of binding to the transcriptional co-activator CBP, with increased levels of phosphorylation enhancing protein binding and function. In contrast, Thr phosphorylation at a single site in p53 (Thr18) functions as an on-off switch in binding to the oncoprotein MDM2.^59,60^ Moreover, the cystic fibrosis transmembrane conductance regulator (CFTR) regulatory region exhibits numerous (mostly Ser) phosphorylation sites that enable a graded (rheostat) response to control interactions with numerous protein targets, with maximum response dependent on multiple phosphorylation events.^61^ In contrast, Thr phosphorylation at two residues of the intrinsically disordered protein 4E-BP2 functions as a regulatory switch, inducing protein folding and changing affinity to binding partners by over 1000-fold.^62^ The data herein provide a context for considering the structural and functional effects of Ser versus Thr phosphorylation: Ser phosphorylation sites inherently exhibit relatively smaller structural effects, suggesting they serve a predominant role when phosphorylation requires a graded (rheostat-like) response, with a full functional event requiring multiple phosphorylation events. In contrast, Thr phosphorylation exhibits significantly more dramatic structural effects, most appropriate for rapid functional effects via step function-like structural switches.

Interestingly, there are a number of proteins that exhibit exceptional specificity for Thr phosphorylation. The protein kinase LKB1, associated with energy homeostasis, diabetes, and organism lifespan, specifically phosphorylates Thr residues in target proteins.^3,63^ FHA domains control protein-protein interactions through the specific binding of proteins phosphorylated on Thr.^4,5^ The pyruvate kinase M2, which acquires protein kinase activity when binding to small molecule regulators, exhibits specificity for phosphorylation at Thr sites.^6,7^ In addition, the protein phosphatase cdc14 selectively dephosphorylates pSer residues, leaving pThr residues.^64^ Moreover, a preference for Ser versus Thr phosphorylation is encoded in protein kinases via the presence of a Phe versus hydrophobic aliphatic residue at the DFG+1 site.^10^ Combined with the observation of differential evolution of Thr and Ser phosphorylation sites,^2^ the data suggest an inherent difference in the native functional application of Ser versus Thr phosphorylation sites. The data herein provide a fundamental structural basis for understanding the differential application of Ser versus Thr phosphorylation within proteins.

Undoubtedly, the results described herein will be context-dependent, with certain sequence and structural contexts capable of promoting dianionic pThr structures that differ from those observed as the predominant conformation of pThr observed herein. Most importantly, monoanionic pThr exhibits only modest conformational preferences, suggesting that local structural contexts that promote monoanionic pThr, or that allow binding of multiple amide hydrogens to pThr, will lead to an alternative structure at pThr.^62^

Interestingly, pThr is a quite sterically hindered amino acid, with both a β-branch and a large phosphate group. Indeed, the steric effects of the phosphate group are observed within pSer, which is commonly observed with an extended side chain conformation that places the sterically demanding phosphate far from the protein backbone (Figure 6, pSer, *χ*_2_ ∼ 180°). However, despite this steric hindrance, pThr nonetheless strongly prefers a highly compact conformation. This result is fundamentally surprising, and indicates the inherent strength of the stereoelectronic effects driving the compact conformation observed herein for dianionic pThr.

Modeling of the effects of protein post-translational modifications is a central goal in order to deduce the complexities of the functional and dynamic proteome. Given the critical roles of protein phosphorylation in intracellular regulation, the development of appropriate force fields for the accurate modeling of the structural effects of post-translational modifications has attracted considerable interest, as well as the recognition that current force fields are insufficient for accurate modeling of the structural effects of phosphorylation.^65-72^ The work herein provides important structural constraints to allow the improved modeling of the effects of protein Ser and Thr phosphorylation.

Intrinsic disorder is an inherent feature of protein phosphorylation sites, with disorder required for protein kinase recognition of substrates.^73,74^ In addition, the majority of phosphorylation sites, including those in protein fragments examined herein, are in regions that remain intrinsically disordered even after phosphorylation. The extant crystallographic data in these proteins thus are dependent on data of protein complexes with these protein segments, which remain quite limited and could potentially be driven by coupled protein binding and folding rather than reflecting the inherent conformational ensemble of these protein segments. The data herein suggest that Thr phosphorylation inherently induces a high degree of local order in proteins, including within intrinsically disordered regions of proteins. In contrast, Ser phosphorylation is capable of inducing a similarly ordered structure, but exhibits a significantly larger range of preferred conformations. These insights have broad potential application in understanding residual and induced structure within disordered regions of proteins.

## Conclusions

We have investigated the inherent structural effects of Ser versus Thr phosphorylation. We have found, through a combination of biophysical analysis in peptides, small molecule X-ray crystallography, analysis of the PDB, and calculations, that Thr phosphorylation induces a particularly strong disorder-to-order conformational change. Dianonic pThr strongly prefers a highly ordered conformation, with a compact *ϕ* ∼ −60°; *ψ* ∼ –30° or +130° (α-helix or PPII); and defined *χ*_1_ (∼ –60°) and *χ*_2_ (∼ +120°). Dianionic pThr has a strong preference for a non-covalently bicyclic structure stabilized by three non-covalent interactions, an intraresidue phosphate-amide hydrogen bond, an interresidue n→π* interaction organizing consecutive carbonyls, and a novel n→σ* interaction between the phosphate Oγ lone pair and the Cβ-Hβ σ*. In combination with the known preference of non-phosphorylated Thr for an extended conformation, these data suggest an inherent difference between phosphorylation of Thr and phosphorylation of Ser residues in proteins, with a greater structural change on Thr phosphorylation than Ser phosphorylation. Thr phosphorylation results in more step-like changes in protein structure and function, while Ser phosphorylation results in more graded or rheostat-like structural and functional changes. These results have broad implications in the differential regulation of proteins by Thr versus Ser phosphorylation.

## Experimental

### Peptide synthesis

Protein-derived peptide sequences are from the indicated residues of the human protein. The peptide sequence for JunD was derived from the murine JunD. Dipeptides were synthesized by solution-phase methods. Larger peptides were synthesized by standard Fmoc solid-phase peptide synthesis, subjected to cleavage from resin and deprotection, and purified to homogeneity via reverse-phase HPLC. Phosphorylated peptides were synthesized via trityl-protected serine or threonine residues, selective trityl deprotection, phosphitylation with O,O’-dibenzyl,N,N-diisopropyl phosphoramidite, and oxidation with *t*-butyl hydroperoxide. Peptides were characterized for identity by mass spectrometry and NMR spectroscopy.

### Circular dichroism

CD spectra were collected on a Jasco J-810 Spectropolarimeter in a 1 mm cell in water containing 5 mM phosphate buffer (pH 8 or as indicated) and 25 mM KF at 25 °C unless otherwise indicated. Data on the Plk1_L_ peptide were collected at 2 °C. Data represent the average of at least three independent trials. Data were background corrected but were not smoothed. Error bars are shown and indicate standard error.

### NMR spectroscopy

NMR spectra were collected on a Brüker 600 MHz spectrometer with a cryoprobe or TXI probe. ^31^P NMR data were collected on a Brüker 400 MHz spectrometer with a BBFO probe. Unless otherwise indicated, NMR experiments were conducted at 298 K in 90% H_2_O/10% D_2_O or 100% D_2_O with 5 mM phosphate buffer (pH 4 for non-phosphorylated peptides, pH 7.5 for phosphorylated peptides) and 25 mM NaCl. 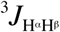 and 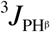 coupling constants were determined in 100% D_2_O to eliminate coupling to the amide hydrogen H^N^.

### PDB analysis

The RCSB protein data bank (PDB) was probed for sequences with threonine, serine, phosphothreonine (3-letter code TPO), and phosphoserine (3-letter code SEP) residues. Serine and threonine were identified from a non-redundant database of known protein structures with ≤ 25% sequence identity and ≤ 1.25 Å resolution using the PISCES server. A total of 14,943 serine and 13,942 threonine residues were identified.^75^ The number of protein structures solved through X-ray crystallography with phosphorylated threonine and serine is significantly lower. To increase the database size for reliable statistics, the conditions for the non-redundant sequence database were relaxed significantly to ≤ 95% overall sequence identity. These structure and sequence requirements led to the identification of 229 phosphothreonine residues from 141 pdb files and 374 phosphoserine residues from 220 pdb files (Tables S2 and S3), which includes identical chains from an individual pdb structure. For those pdb files with multiple sequence-equivalent chains, only one chain (among the sequence-equivalent chains) was included for analysis (either the first chain in the file, or the most ordered chain if differences in order were identified) and final plots, such that it represents an unbiased analysis of phosphorylated residues from all the different protein structures without repetitive sequences. The database was finally curated by rejecting structures with > 2.5 Å resolution. For accurate representation of structural elements, any structure with missing residues immediately preceding or following a phosphorylated residue, or phosphorylated residues missing any side chain heavy atom (either due to inherent disorder or to limitations of the X-ray structure) was excluded from the analysis. This process led to the identification of 155 phosphothreonine residues and 234 phosphoserine residues included in the final statistical plots (Figure 5, Tables S2 and S3). The dihedral angles were calculated using in-house programs written in the Perl programming language.

### X-ray crystallography

#### Racemic crystallization

A 1:1 mixture of l- and d-peptide enantiomers of 4-iodo-benzoyl-phosphothreonine-proline-NH_2_ (pH adjusted to 7.8 in water and dried) was dissolved in 1:1 MeOH:*t*-BuOH and subjected to crystallization at 298 K by sitting drop vapor diffusion using a *t-*BuOH reservoir. The racemic peptide crystallized over a period of one week. The crystals were diffracted using Cu radiation at 30 seconds per frame. Disorder in the crystals, potentially due to the large volume occupied by water (10 water molecules localized per asymmetric unit) and sodium atoms that are expected to be relatively mobile, resulted in a maximum resolution of 0.87 Å. No symmetry higher than triclinic was observed, and refinement in the centrosymmetric space group P–1 yielded chemically reasonable and computationally stable results in refinement.^76^ Two similar, yet symmetry-unique, molecules of the peptide were observed in the asymmetric unit. Only three sodium atoms can be localized per two peptides; it is likely that the fourth expected counterion resided in a highly disordered portion of the electron-density map centered near the origin (469 Å^2^, 122 e) that could not be modeled, and that was thus treated as diffused contributions.^77^ *4-Iodobenzoyl-phospho-*l*-threonine-*l*-proline amide* The single-enantiomer compound was subjected to crystallization via vapor diffusion of a solution of 50% methanol in *tert*-butanol (v/v) with *tert*-butanol at room temperature. Crystals formed over a period of two years. The structure was solved by standard methods in X-ray crystallography, with a resultant resolution of 0.80 Å. For this crystal structure, all non-hydrogen atoms and the hydrogen atoms involved in hydrogen bonding were localized from the difference map. These structures were deposited in the Cambridge Structural Database under the depositary numbers CCDC 1877191 and 1877192.

### Calculations

Calculations were conducted using Gaussian09^78^ on a series of models (Ac-pThr-NMe_2_, Ac-pThr-NMe_2_•bridging H_2_O, Ac-pThr-Pro-NMe_2_, Ac-pThr-Pro-NMe_2_•bridging H_2_O, Ac-pSer-NMe_2_, Ac-pSer-NMe_2_•H_2_O, Ac-pSer-Pro-NMe_2_) that were initially derived from crystallographically observed structures. Structures were chosen to include an acetyl group at the N-terminus, which provides an appropriate balance between computational simplicity and appropriately representing the electronic properties of the donor carbonyl in an n→π* interaction.^79^ Molecules were analyzed with a dimethyl amide on the C-terminus in order to avoid potential artefacts due to the presence of an unsatisfied hydrogen bond donor at the N-terminus. The charge and multiplicity of the system were set at –2 and 1, respectively. Initial geometry optimization calculations were conducted in Gaussian using the M06-2X DFT method^80^ and the 6-311++G(d,p) basis set, using implicit solvation with water via a continuum polarization model (IEFPCM). These initial models were then subjected to further optimization with the 6-311++G(2d,2p), then 6-311++G(3d,3p) basis sets. The resultant models were then analyzed using frequency calculations. If an imaginary frequency was observed, the models were subjected to further geometry optimization until a final structure was obtained that had no imaginary frequencies. All final geometry-optimized structures had zero imaginary frequencies.

Natural bond orbital (NBO) analysis^81^ of the geometry-optimized Ac-pThr-Nme_2_ structure at M06-2X level of theory with the 6-311+G(3d,3p) basis set was conducted post-optimization to provide insights into the origins of the observed conformational preferences. The NBO method transforms a calculated wavefunction into localized form, which is used in quantum chemistry to calculate the distribution of electron density in atoms and in bonds between atoms. The stabilization energies via eclipsing C-H/O-P bonds, n→π*, and phosphate-amide hydrogen bond interactions of the geometry-optimized structures were calculated by using second-order perturbation theory as implemented Gaussian09. The relevant NBO orbitals that are involved in the three major non-covalent interactions in this study were visualized using GaussView 5 with isovalues of 0.02.

Structures of the minimal molecules isopropyl phosphate and ethyl phosphate were analyzed via geometry optimization and via scans of the HCOP torsion angle (equivalent to a scan of the χ_2_ torsion angle in phosphothreonine or phosphoserine, respectively), in order to understand the side chain conformational preferences of phosphothreonine *versus* phosphoserine. Geometry optimization was conducted at the MP2 level of theory with the 6-311++G(3d,3p) basis set in implicit water, in the dianionic, monoanionic, and neutral forms of phosphate. Geometry-optimized structures were analyzed by frequency calculations, with no imaginary frequencies observed. These structures were then subjected to HCOP torsion angle scans, using the MP2 method and the 6-311++G(2d,2p) basis set in implicit water.

## Supporting information

cif file of crystal structure

cif file of crystal structure

## Acknowledgements

We thank NIH (GM93225), NSF (BIO-1616490), and the University of Delaware for funding. We thank Brian Kuhlman (UNC-Chapel Hill) for initial help with PDB analysis. We thank Alaina Brown, Krista Thomas, and Agata Bielska for preliminary CD experiments on JunD and Gsn1.

