## Supplementary figures legends for "An Inherent Structural Difference Between Serine and Threonine Phosphorylation: Phosphothreonine Prefers an Ordered, Compact, Cyclic Conformation"

**Supplementary Tables**

**Table S1**. Average values of NMR-derived data at 298 K for all peptides in Figure 1. Data for monoanionic phosphorylated peptides were collected at pH 4. Data for dianionic phosphorylated peptides were collected at pH 7.5-8.0. Changes indicate change in the indicated value from the non-phosphorylated residue to the dianionic phosphorylated residue.

|  | S  Ser(OH) | pS^–^Ser(OPO_3_H^–^) | pS^2–^Ser(OPO_3_^2–^) | T  Thr(OH) | pT^–^  Thr(OPO_3_H^–^) | pT^2–^Thr(OPO_3_^2–^) | change,  S to pS^2–^ | change,  T to pT^2–^ |
| --- | --- | --- | --- | --- | --- | --- | --- | --- |
| H^N^ δ, ppm | 8.37 | 8.52 | 8.87 | 8.25 | 8.44 | 9.51 | +0.50 | +1.23 |
| N^H^ δ, ppm | 118.1 | 117.3 | 119.5 | 118.4 | 119.4 | 124.0 | +1.4 | +5.6 |
| ^3^*J*α_N_, Hz | 6.8 | 6.4 | 5.6 | 7.3 | 6.5 | 3.5 | –1.2 | –3.8 |
| Hα δ, ppm | 4.65 | 4.82 | 4.68 | 4.55 | 4.60 | 4.34 | +0.03 | –0.21 |
| Cα δ, ppm | 53.4 | n.d. | 53.0 | 57.1 | n.d. | 58.5 | –0.4 | +1.4 |

**Table S2**. Proteins containing phosphothreonine and structural data used in Figure 5.

**Table S3**. Proteins containing phosphoserine and structural data used in Figure 5.

**Table S4**. Results of geometry optimization calculations, with comparisons to small-molecule data from X-ray crystallography.*^a^*

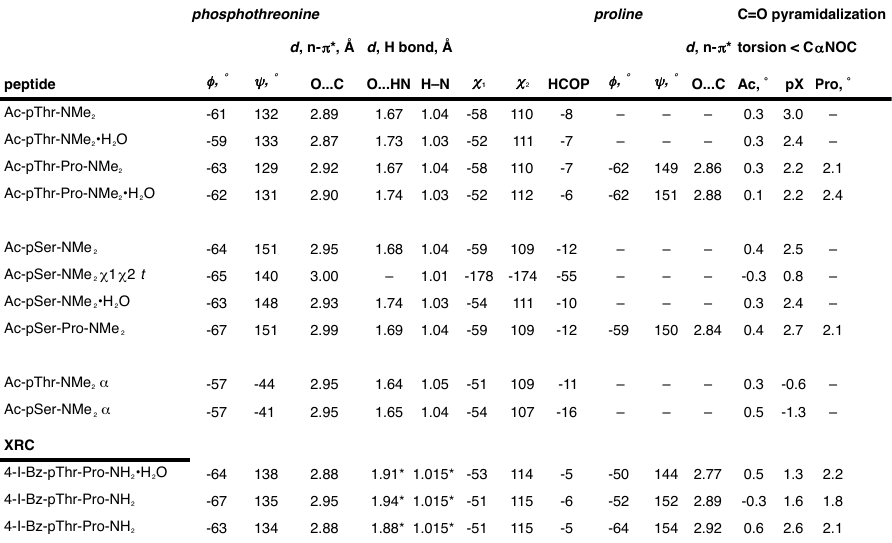

*^a^* Calculations were conducted using the M06-2X DFT method and the 6-311++G(3d,3p) basis set in implicit water. Structures with •H_2_O appended to the end of the peptide name indicate structures with a bridging water molecule, as was observed in the single-enantiomer crystal structure. The torsion angles in the structure of the D-peptide determined by X-ray crystallography were converted to their mirror-image values to simplify comparison across peptides. * These numbers are based on an H–N bond length normalized to 1.015 Å.
